# SSPSPredictor: A Sequence and Structure based Deep Learning Model for Predicting Phase-Separating Proteins

**DOI:** 10.64898/2026.03.30.715224

**Authors:** Tinglan Wang, Shaofeng Liao, Yifei Qi, Zhuqing Zhang

## Abstract

Liquid-liquid phase separation (LLPS) underlies the formation of biomolecular liquid condensates (also referred to membraneless organelles, MLOs), which are essential for spatially organizing various biochemical processes within cells. Proteins that play a key role in driving condensates formation are termed phase-separating proteins (PSPs). Given experimental identification of PSPs remains labor-intensive and time-consuming, multiple computational tools have been developed based on empirical features or deep learning. In this study, we propose SSPSPredictor, a novel multimodal predictive model for PSPs with folded or intrinsically disordered structures, leveraging the fusion of sequence information from a protein language model ESM-2 and structural insights from a graph neural network GVP. Compared with existing tools, SSPSPredictor achieves balanced performance in identifying endogenous PSPs, predicting relative LLPS propensities, and recognizing key regions that drive LLPS. Moreover, SSPSPredictor exhibits good interpretability in identifying driving regions along protein sequences, although no relevant supervision was provided during training. Further predictive analysis of the human proteome using SSPSPredictor reveals that the proportion of intrinsically disordered proteins (IDPs) undergoing LLPS is significantly higher than that of folded proteins. In addition, pathogenic variants, especially those located in disordered regions, exhibit higher LLPS propensity than other mutations, uncovering a link between LLPS and diseases at the amino acid level.

## INTRODUCTION

Biomacromolecules within cells are generally non-uniformly distributed, forming distinct compartments that can be broadly categorized into membrane-bound organelles and membraneless organelles (MLOs). Many MLOs are dynamically stable liquid condensates containing numerous proteins and nucleic acids^1,2^, which are usually involved in various biological processes, including signal transduction, immune synapse formation, nuclear transcription, etc.^3^. These biomolecular condensates are widly deemed to be formed through liquid-liquid phase separation (LLPS)^4,5^, which occurs when the concentration of biomolecules surpass critical threshold under certain condition, and exhibits the coexisting of two distinct liquid phases: a concentrated phase (liquid condensates) and a dilute phase^5,6^. Proteins primarily driving condensates formation are known as scaffolds in LLPS, and also coined as phase-separating proteins (PSPs). Extensive experiments have been demonstrated that many intrinsically disordered proteins/regions (IDPs/IDRs) play central roles in LLPS, because they can provide flexibility and multivalent interaction sites which are crucial in liquid condensates formation. A number of IDPs or proteins containing IDR(s) have been discovered belonging to PSPs. Meantime, some proteins with tandem repeatedly folded domains have been found to undergo LLPS as the domain-domain interaction providing the multivalent interaction sites. Accurate identification of PSPs is important for elucidating physiological and pathological mechanisms of biomolecular condensates.

Given that experimental identification of PSPs remains labor-intensive and time-consuming, multiple computational tools have been developed^7–9^. Among these algorithms, some of them utilize engineered features derived from prior knowledge or statistical enrichment analyses to build the model, such as Pscore based on planar sp2 π-π interactions^10^, FuzDrop based on the entropy differences between bound and unbound states^11^, as well as LLPhyScore^12^, PhaSePred^13^ etc. based on multiple statistical features. Other predictors such as PSPredictor^14^, DeePhase^15^ and PredLLPS_PSSM^16^ leverage word2vec or its modification to learn protein sequence latent features. Recently, a couple of tools were built on pretrained protein language models ESM-2^17^, which has been demonstrated to have strong capacity to understand semantic of protein sequences. Its embedding application has achieved excellent performances on multiple related task, including phase transition prediction. However, most of the above predictors focus on PSPs dominated by largely disordered proteins, and some even use folded domains as negative training set, which may lead to bias results. To address this limitation, Hu et al.^18^integrates structural information from AlphaFold2 with residue-level features of IDPs, build the model PSPire based on machine learning, and showed notable improvement in identifying PSPs without IDRs. This approach highlights the importance of leveraging three-dimensional structural data for PSPs prediction.

In this study, we present a novel algorithm for identifying PSPs which incorporates protein sequence information based on ESM-2 and structural information generated by AlphaFold2^19^. We tried one graph neural networks (GNNs) for protein structure presentation, one is GVP^20^ and the other is SPIN^21^. GNNs have shown great promise in protein-related tasks such as structure-based property prediction^22^ and interaction modeling^23,24^. In addition, the parallel and sequential integration manners of ESM-2 and GNNs were discussed, and the module of attentional mechanism^25^ was adopted which endows the predictor interpretability. After a series of validations on external test sets, the final model SSPSPredictor exhibit a balanced and overall excellent performance. This robust tool for PSPs prediction would facilitate deeper researches in biomolecular phase separation.

## MATERIALS AND METHODS

### Workflow

We incorporated both protein sequence and structure in model construction. For protein sequence information, the widely used pretrained protein language model ESM-2 was applied to extract contextual sequence information. It generates residue-level embeddings that capture semantic and evolutionary features through masked language modeling (MLM)^26^. Specifically, the variant “esm2_t33_650M_UR50D” was used in this study, which was trained on the “UR50/D 2021 04” dataset, producing 1280-dimensional vectors for each residue. For structural features, predicted conformations from AlphaFold2 were used as input. Meanwhile, two GNN-based structure encoders, GVP and SPIN-CGNN, were applied through which proteins were represented as graphs, where each residue is treated as a node (V) and their interactions are treated as edges (E). GVP excels at integrating scalar and vector features within a global coordinate framework, and its core advantage lies in strict SE(3) equivariance, which ensures consistent feature representations under arbitrary 3D rotations and translations of the input data. It processes input information primarily by separating scalar and vector channels. In protein structures, node feature information (scalars) is accompanied by geometric information such as interatomic directions and distances (vectors) between nodes. GVP layers are capable of simultaneously capturing and processing both types of information, thereby avoiding loss of geometric information inherent in conventional GNNs when handling spatial data. SPIN-CGNN enhances the performance of fixed-backbone protein sequence design by improving graph construction and edge update mechanisms. The model employs a contact-map-based graph construction method, dynamically determining connections between nodes through distance thresholds to effectively handle differences of local structural density. Meanwhile, SPIN-CGNN introduces a Second-Order Edges (SOE) update strategy to enhance message-passing efficiency between graph nodes. It also incorporates a Selective Kernel (SK) module to adaptively fuse multi-scale features, optimizing the model’s ability to capture complex structural characteristics. Previous studies have shown that SPIN-CGNN significantly outperforms some state-of-the-art methods, such as ProteinMPNN^27^, across multiple evaluation metrics.

The integration of protein sequence information (derived from ESM-2) and structure information derived from GNNs (GVP or SPIN-CGNN) can be parallel or sequential fusion. We tried the two manners in this study.

Recently, based on ESM-2 architecture and Foldseek^28^ to encode three dimensional structural information, a pretrained model SaProt^29^ was developed, which generates residue tokens representing amino acid types and structural tokens capturing spatial features. We used SaProt (“SaProt 650M PDB” variant, corresponding to ESM-2 model above) to produce 1280-dimensional vectors for each residue for model constructions, to compare with strategies mentioned above.

In addition, attention pooling layers were used to obtain the weighted score for each residue, and a multilayer perceptron (MLP) module was used to output prediction scores and identify PSPs. The attention pooling layers can also identify key residues, ensuring that critical information is preserved for downstream tasks, which provides interpretability for the models. To further enhance the diversity of features captured, we included a structured attention penalization term, which encourages attention heads to focus on specific sequence regions. This penalization item is defined as

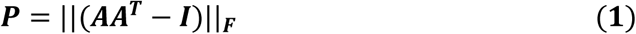

where F represents the Frobenius norm of a matrix, and A denotes the attention matrix, and I is the identity matrix, which acts as a regularization component to ensure robustness and effectiveness of the model.

Overall, six models with different combinations of the components described above were constructed, including ESM_only, SaProt, ESM_GVP_p, ESM_SPIN_p, ESM_GVP_s and ESM_SPIN_s, as the architecture schematics shown in Figure 1. Among them, ESM_only uses the embedding obtained from ESM-2 and does not incorporate structural information, and other five introduced structural information in different ways, where “_p” and “_s” denote parallel and sequential integration strategies, respectively.

**Figure 1.**
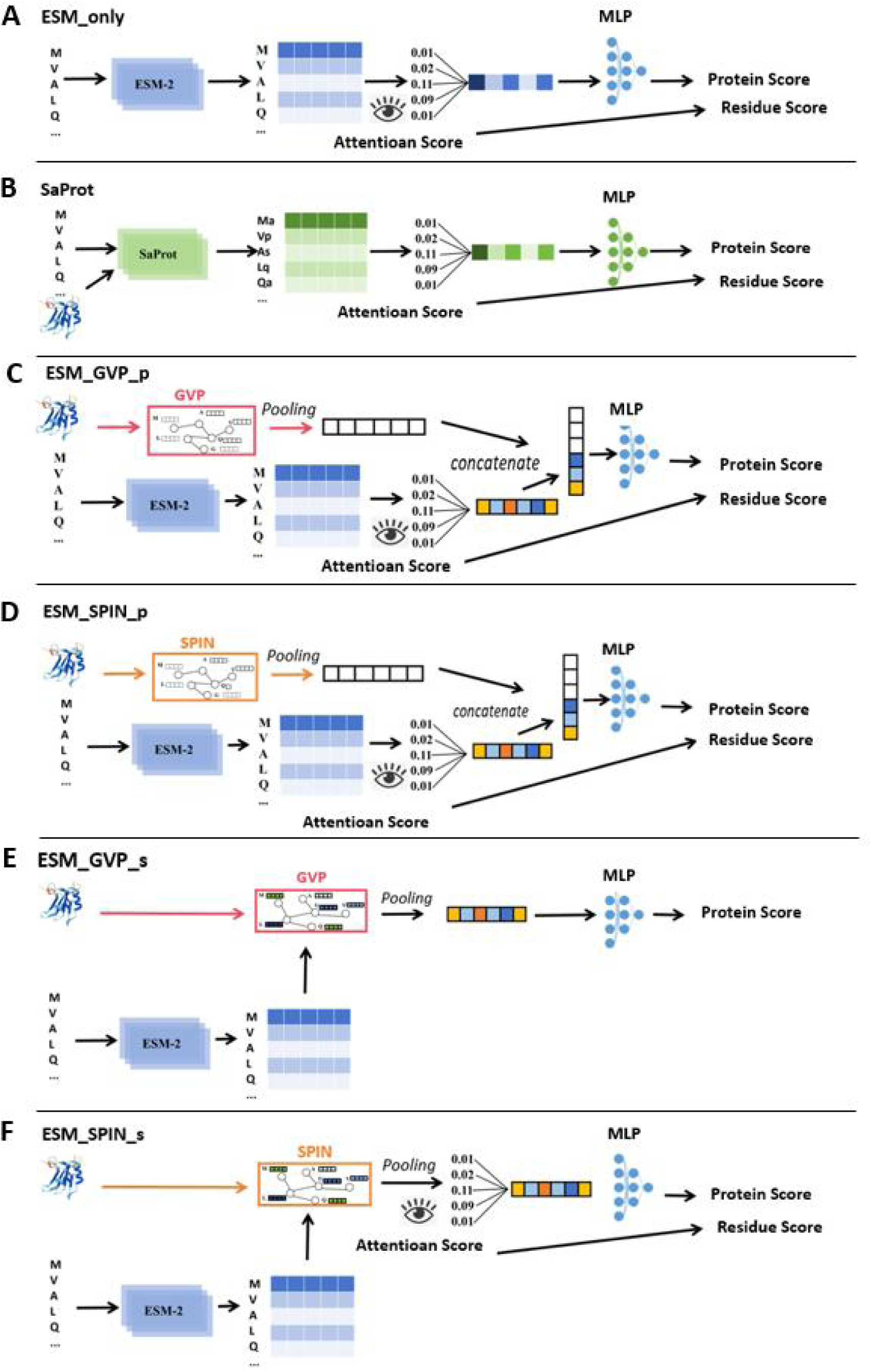
Model Architecture. A: Architecture of the ESM_only model; B: Architecture of the SaProt model. C: Model architecture based on parallel (p) fusion of ESM2 and GVP. D: Model architecture based on sequential (s) fusion of ESM2 and SPIN-CGNN. E: Model architecture based on sequential fusion of ESM2 and GVP. Since GVP employs average pooling to aggregate local features into a global representation, it cannot generate residue-specific scores. Thus, ESM_GVP_s does not output amino acid-level predictions. F: Model architecture based on sequential fusion of ESM2 and SPIN.

### Datasets

We utilized the updated PhaSepDB v2^30^, LLPSDB v2^31^, as well as AlphaFoldDB^32^ to get the datasets for training and test. For the positive dataset, we selected proteins labeled as “PS-self” from PhaSepDB v2 and proteins undergoing LLPS in single-protein systems from LLPSDB v2, indicating that these proteins are capable of undergoing self-assembly via phase separation. Sequences from hetero-oligomers or containing non-natural amino acids, as well as those with fewer than 16 or more than 2048 amino acids, were excluded. To reduce redundancy, we applied the CD-HIT algorithm^33^ with a sequence similarity cutoff of 0.4 and selected one representative sequence per cluster. This resulted in a final positive dataset including 352 sequences. For the negative dataset, we selected sequences from the human proteome in AlphaFoldDB, excluding those present in LLPSDB v2 or PhaSepDB v2, those containing non-standard amino acids, and those not matching the length range of the positive dataset. A total of 1700 sequences (about five times of sequences in the positive dataset) were randomly sampled to construct the negative dataset. To enable fair comparison with existing prediction methods, we used 104 newly updated sequences in the positive dataset as the positive test set, which had not been used to train any of the compared models. The remaining 248 sequences in positive dataset were used as the positive training set. The train-validation dataset consisted of 248 positive and around 248 × 5 negative samples. The positive samples were split into training and validation sets in a 4:1 ratio. The negative training set was obtained by randomly selecting sequences from the negative dataset at five times the number of positive samples, and the remaining sequences were used as the negative test set. The positive test set and the negative test set constitute a “Test set 0” for evaluation and comparison with other models.

In addition, we collected data from references and built several other datasets to evaluate our models, which is briefly described as following,

Test set 1: This dataset is constructed to evaluate model ability to identify endogenously expressed PSPs. Li et al. identified a set of endogenously expressed PSPs through high-throughput experiments^34^. They reported a total of 1518 PSPs, among which 538 sequences had never been reported before. We removed the sequences that do not meet the length restriction (<50, to meet the requirements of the model PredLLPS_PSSM for comparison) or contained non-natural amino acids from the 538 proteins, and obtained a total of 524 sequences.

Test set 2: This dataset was constructed to evaluate whether models could predict relative LLPS ability based on the saturated LLPS concentration for a group of mutations. Bremer et al.^35^ conducted extensive mutation experiments using the prion-like low-complexity domain (PLCD) from isoform A of human hnRNPA1 (referred to as A1-LCD) as a model protein, to investigate the impact of different types of amino acids on the propensity for protein LLPS. The saturation concentrations of various mutated proteins were measured, above which condensates can form under the same conditions, including pH, salt, concentration and temperature. This data set provides a valuable criterion for evaluating performance of PSPs predictors in LLPS propensity prediction. Same as the Ref 8 from Bremer et al.’s experiments, we selected two groups of mutant sequences based on the experimental temperature. One group includes 17 sequences with mutations primarily focused on aromatic or charged residues, and the saturation concentrations were measured at 4°C (A1LCD_4). The other group includes 13 sequences that specifically focus on the role of glycine (Gly) and serine (Ser) residues, and the saturation concentrations were measured at 20°C (A1LCD_20).

Test set 3: This dataset includes the driving regions contained in PSPs collected from the database PhaSePro^36^, which includes 121 proteins with phase-separating regions (driving regions) validated by experiments. It was used to evaluate the model ability to identify key residues in PSPs. Each residue in the dataset is simply binarized (0/1 for presenting the residue is key one or not) for comparison of models in convenient way.

Test set 4: This dataset was derived from ClinVar^37^, which contains 126,899 variants from the human proteome, including 82,643 benign and 44,256 pathogenic variants. These variants are used to analyze the relation of pathogenic mutations with LLPS propensity in this study.

### Model training and evaluation

Cross-training resulted in five architectures. Each was trained on a distinct negative subset (all negative samples were evenly split into five sets) to enhance data diversity, yielding a total of five sub-models. The Optuna framework38 was used for systematic hyperparameter optimization. We transformed the binary classification of PSPs into a regression problem, training the model to output a score representing the LLPS propensity of a protein. The score was subsequently binarized to derive the final class label. Finally, predictions from the five sub-models were integrated – either by averaging or by majority voting, depending on the task – to produce the final outputs, including residue-level scores, protein-level scores, and binary classification labels.

The models were evaluated using several standard metrics: the area under the receiver operating characteristic curve (AUROC), the area under the precision–recall curve (AUPRC), F1-score, accuracy, false positive rate (FPR), and false negative rate (FNR). To calculate F1-score, accuracy, FPR, and FNR, prediction scores were converted into binary labels using the threshold corresponding to the point nearest the top-left corner of the ROC curve. Their formulas are given as follows:

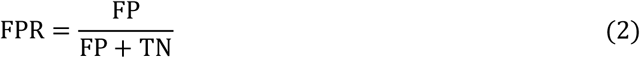

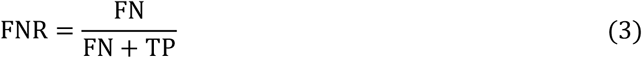

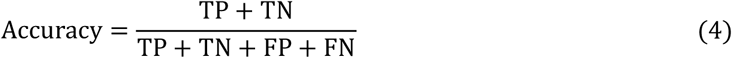

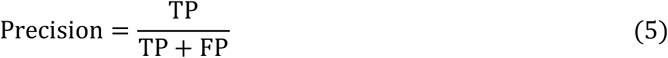

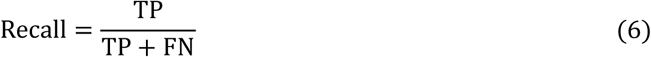

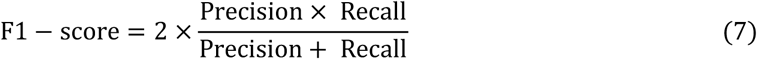

In addition, we used the Spearman correlation coefficient (ρ) to evaluate the correlation between two sets of variables:

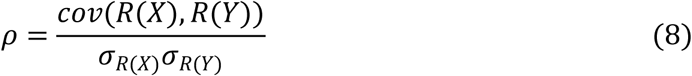

Here, the numerator represents the covariance of the rank variables, and the denominator represents the product of their standard deviations. The parameter ρ ranges from −1 to +1, where 0 indicates no correlation, and positive and negative values indicate positive and negative correlations between two variables, respectively.

## RESULTS

### The performance in identifying PSPs

We compared the six models (using the average value of five sub-models for each model) in this study with several published predictors of PSPs, including DeePhase^15^, PSPredictor^14^, FuzDrop^11^, LLPhyscore^12^ and PLAAC^39^. These previous models have demonstrated outstanding predictive performance^8^. Among the models trained by LLPhyscore on different datasets, we selected the model trained on the human proteome for comparison. Additionally, recently developed tools-PSPHunter^40^, PSPire^18^, ParSeV2.0^41^ and PredLLPS_PSSM^16^ were not trained on the aforementioned data and were thus also included in the comparison. We compared the prediction performance of these algorithms with the six models we constructed based on the Test Set 0 (see the above section “Datasets”), as shown in Figure 2A. When the models ranked by AUROC, six of the top seven models were ESM2-based from this study. The remaining model is PSPire, which incorporates structural information and ranks third, exhibiting a relatively lower AUPRC but a slightly higher F1-score compared to the six ESM2-based models. As a whole, according to AUROC, the performance of SaProt surpasses other models, followed by ESM2_only, indicating the efficacy of incorporating structural information in enhancing model performance. Moreover, the GVP-based model outperforms the SPIN-based model. One possible explanation for this phenomenon is that SPIN possesses a larger number of parameters compared to GVP, making it prone to overfitting on small datasets and resulting in suboptimal performance. We then examined whether these models are capable of identifying endogenous PSPs in Test Set 1. This dataset comprises 524 endogenous PSPs that were previously unreported and thus excluded from the training data of all models. Figure 2B shows the identification performance on this dataset. PredLLPS_PSSM model stands out by identifying 252 PSPs, much more than other models. DeePhase, which is based on the word2vec language model and sequence features, identifies 209 PSPs. Next are ESM_GVP_p, ESM_GVP_s, and PSPire, each identifying more than 150 PSPs. The number of overlapping PSPs predicted by any pair of the models is visualized in Figure S1. The largest overlap occurs between PredLLPS_PSSM and DeePhase, followed by that between ESM_GVP_p and ESM_GVP_s. Notably, all algorithms perform suboptimally on Tset Set1 (identification rate < 50%), suggesting that their training data lack sufficient diversity.

**Figure 2.**
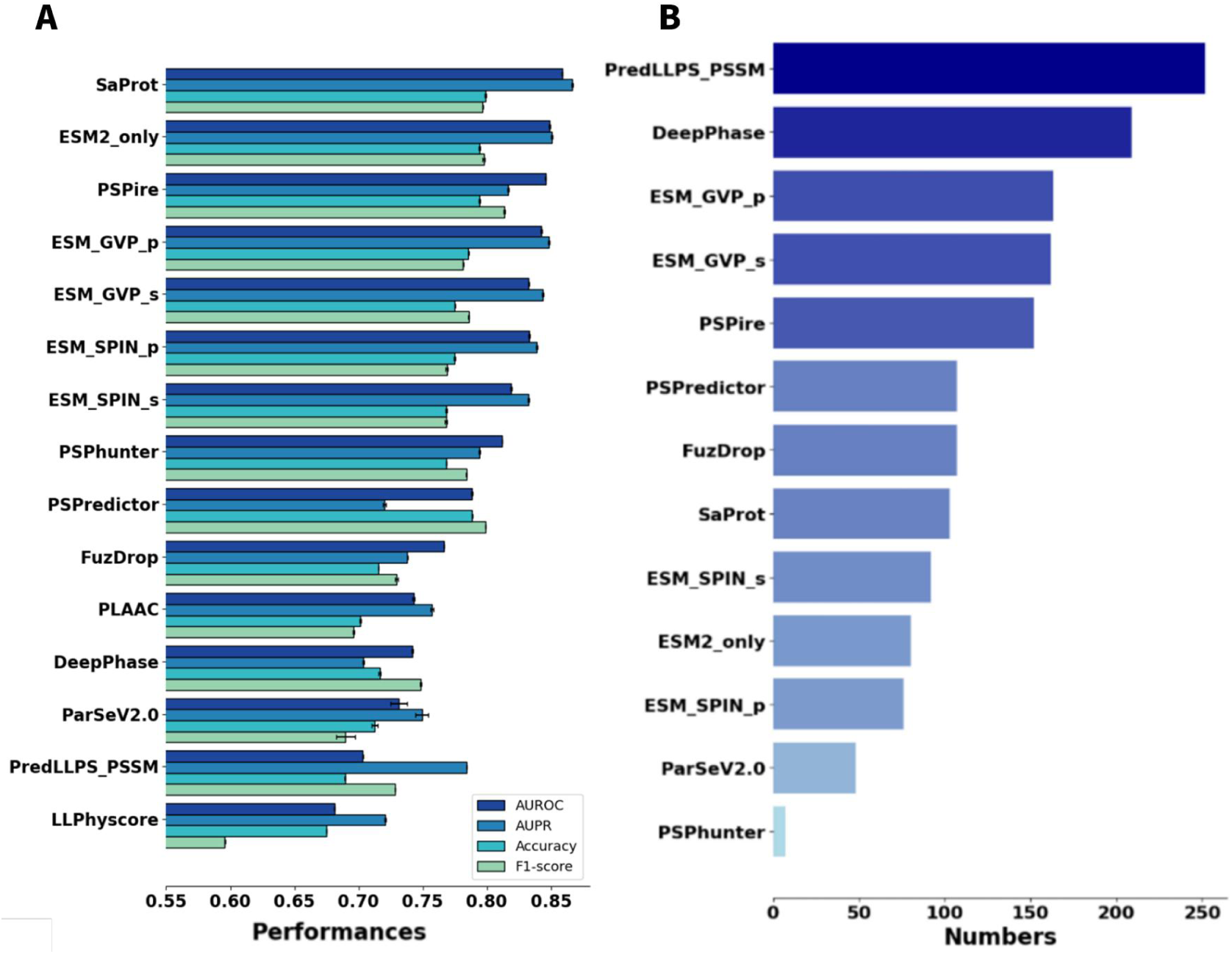
Evaluation on Test set 0 and Test set 1. A: Four evaluation metrics (AURCO, AUPR, Accuracy, and F1-score) of different models (For all our models, the result is the average of the scores from the five sub-models.) on Test set 0; B: The number of endogenous phase-separating proteins identified by different models on Test set 1. For all our models, the result is determined by the voting of the five sub-models.

### The performance in prediction of LLPS propensity

Using Tset Set2—which contains experimental saturation concentrations for mutants of the prion-like low-complexity domain (PLCD) of human hnRNPA1 isoform A (hereafter A1LCD)^35^—we compared the LLPS propensities predicted by our models with those of other methods. The dataset includes extensive saturation concentration of phase separation for extensive mutation sequences, with one group at 20°C (13 sequences) and the other at 4°C (17 sequences). Because a lower saturation concentration indicates a greater propensity for phase separation, a more negative correlation coefficient between saturation concentration and a model’s prediction score reflects better performance. As shown in Figure 3, most models show better correlation on A1LCD_20 (red bars) than on A1LCD_4 (blue bars). This may be because 20°C is closer to room temperature, at which most training data were collected. Specifically, ESM_only, ESM_GVP_p, and ParSeV2.0 perform best on A1LCD_20, whereas ESM_only, ESM_GVP_p, and ESM_GVP_s rank in the top three on A1LCD_4. The poor performance of other tools suggests they cannot reliably predict LLPS propensity for A1LCD mutants.

**Figure 3.**
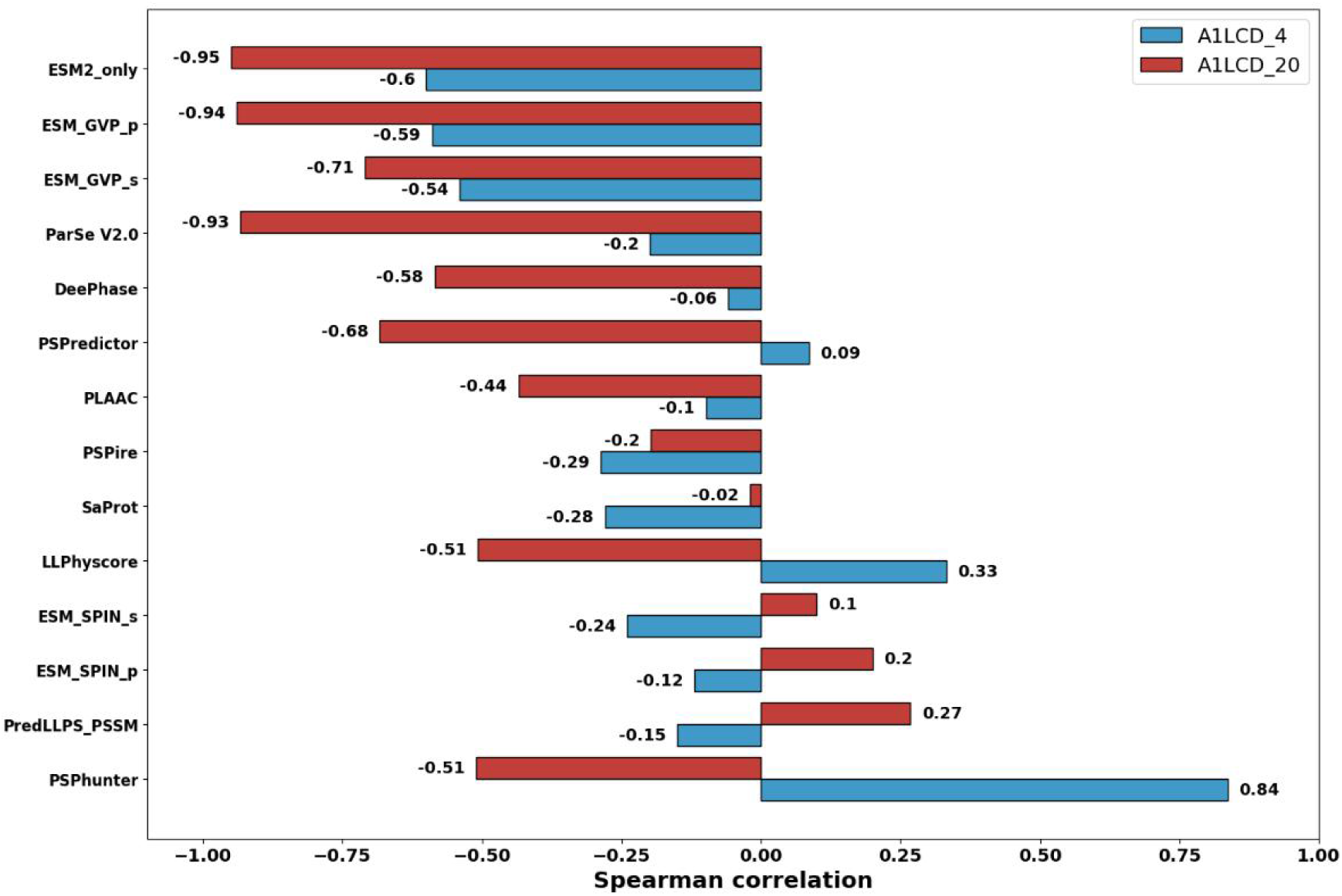
Evaluation on Test set 2. Spearman correlation coefficients were calculated between the predicted scores of different prediction algorithms and the saturation concentrations. The models are sorted from top to bottom in ascending order of the mean value. For all our models, the result is the average of the scores from the five sub-models.

### Attention score interprets crucial residue of proteins in LLPS

The LLPS process of a phase-separating protein (PSP) is typically driven by specific crucial residues or regions within its sequence. In our models, a residue-level importance score for each predicted PSP was derived by scaling the normalized attention-pooling weights by the protein’s sequence length. These scores were averaged across the five submodels and evaluated against experimentally validated phase-separating regions. Test Set3 comprises 121 PSPs from PhasePro, with each residue labeled as 1 if it lies within a phase-separating region and 0 otherwise. Treating the 86,660 residues from the 121 PSPs in Test Set3 as a binary classification task, we first compared the four models developed in this study. As shown in Figure S2, both ROC and PRC curves indicate that the ESM_GVP_p model outperforms the other four models. Figure S3 reveals a clear separation between the distributions of residue scores inside versus outside phase-separating regions in Test Set3, with residues within driving regions exhibiting significantly higher scores. This suggests that integrating ESM2-based sequence embeddings with GVP-derived structural representations can effectively capture residue-level features critical for LLPS, despite the absence of residue-level labels during training.

We then compared the performance of ESM_GVP_p with that of other existing predictors. Among existing tools, FuzDrop, PSPHunter, and LLPhyScore provide residue-level scores and can identify key amino acids. The ROC and PRC shown in Figure 4 demonstrate that ESM_GVP_p outperforms the three methods mentioned in identifying crucial residues in PSPs. Meanwhile, by applying a threshold derived from the point closest to the top-left corner of the ROC curve, we extended the evaluation to additional methods—including PSPire, ParSeV2.0, and PLAAC—that do not output continuous residue scores but can be assessed via binary predictions. Given that residue importance is context-dependent, we computed per-protein accuracy and F1-scores for all 121 PSPs in Test Set3. As shown in Figure 5, ESM_GVP_p achieves the highest accuracy and F1-score distributions across the 121 PSPs, outperforming other methods. FuzDrop and LLPhyscore rank second and third, respectively; notably, FuzDrop was trained on this same dataset.

**Figure 4.**
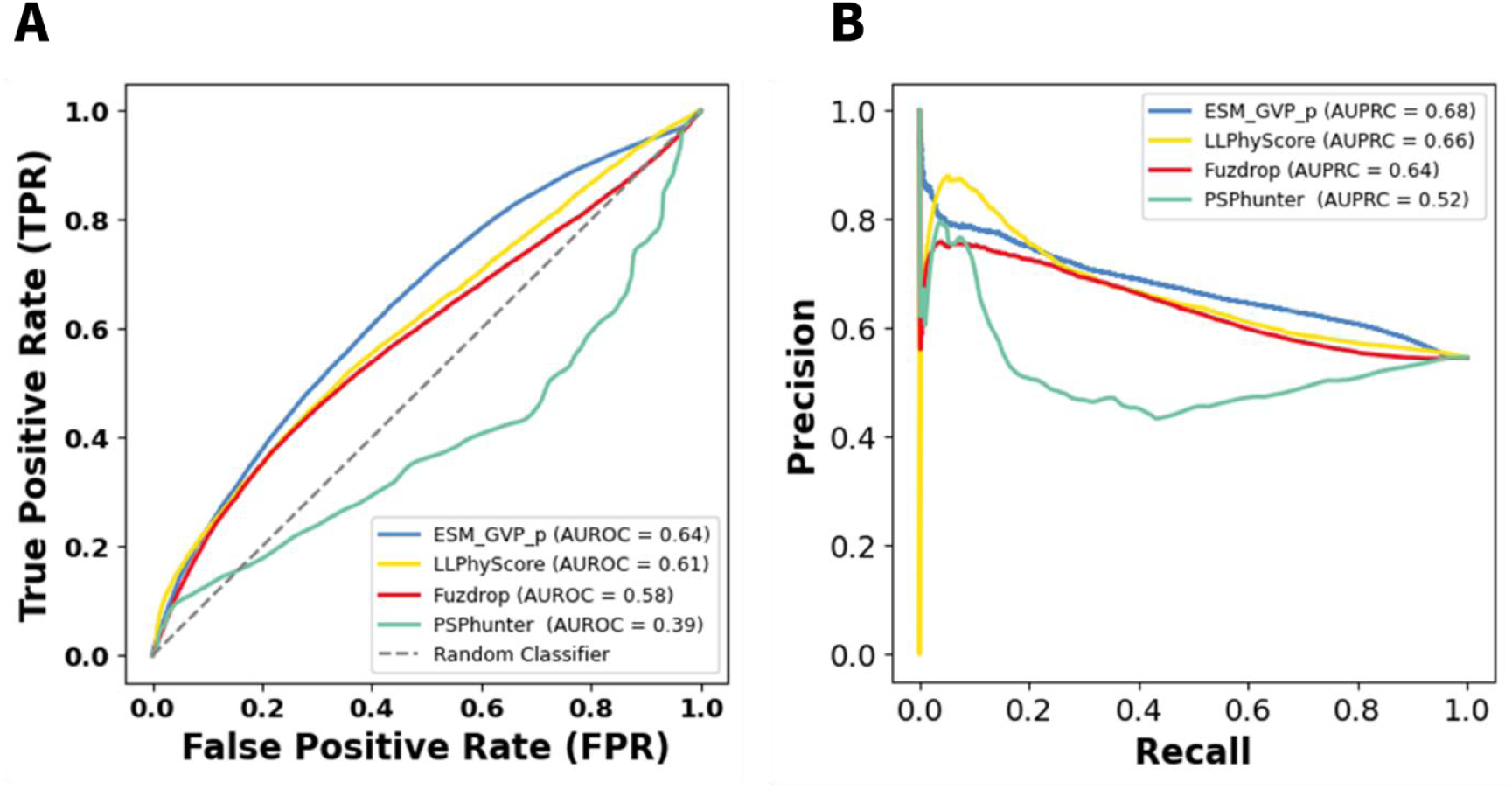
Evaluation on Test set 3. ROC curves (A) and PR curves (B) are obtained across all amino acids of the dataset. Amino acids that is experimentally validated within phase separation-prone regions were set as 1 and other vise as 0. For all our models, the result is the average of the residual scores from the five sub-models.

**Figure 5.**
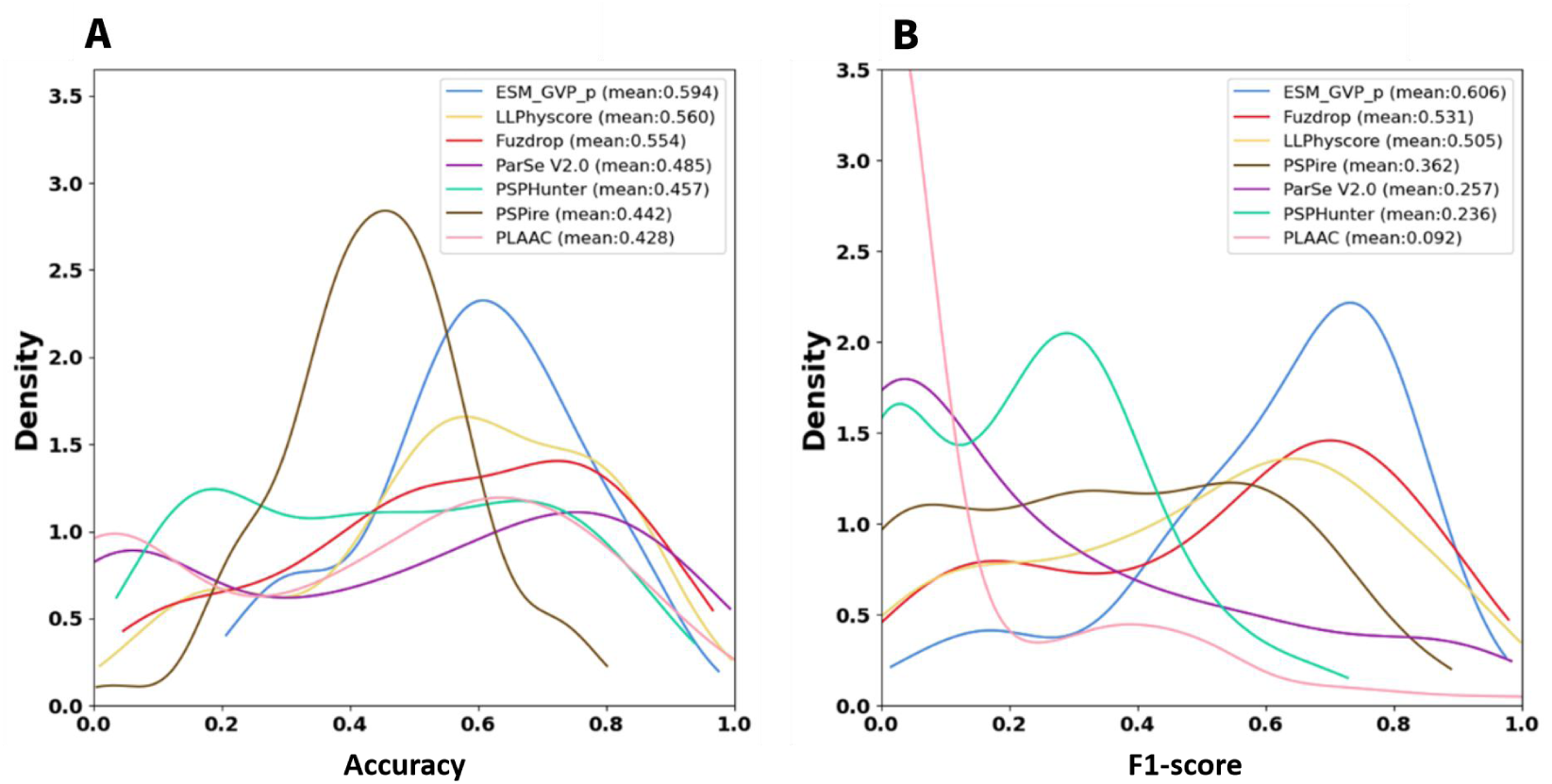
Accuracy and F1-score of different models in identifying the phase-separating driving regions of proteins in Test set 3. For all models involved in Figure 4, the threshold is selected as the one corresponding to the optimal point on the ROC curve.

We further selected nine well-studied PSPs that have attracted widespread attention and visualized their experimental validated and predicted phase-separating regions along sequences. As shown in Figure 6, black bars denote experimentally verified driving regions, while colored tracks (from top to bottom) represent predictions from ParSeV2.0, FuzDrop, PSPHunter, PSPire, PLAAC, LLPhyscore, and SSPSPredictor, respectively (with predicted crucial residues marked by short vertical bars). This figure shows that SSPSPredictor can identify most of the phase-separating segments in these nine proteins, while PSPire and FuzDrop often yield opposing predictions: the former emphasizes structured regions, whereas the latter prioritizes intrinsically disordered regions. For instance, the experimentally defined phase-separating region of Tau spans residues 244–368. AlphaFold2 predicts this segment to be largely ordered (high pLDDT). FuzDrop does not predict this segment as a driving region, while PSPire and SSPSPredictor do. Conversely, for the C-terminal domain of TDP-43 (residues 263–414), the pattern is reversed. That is, PSPire does not predict this segment as a driving region, while FuzDrop and SSPSPredictor do.

**Figure 6.**
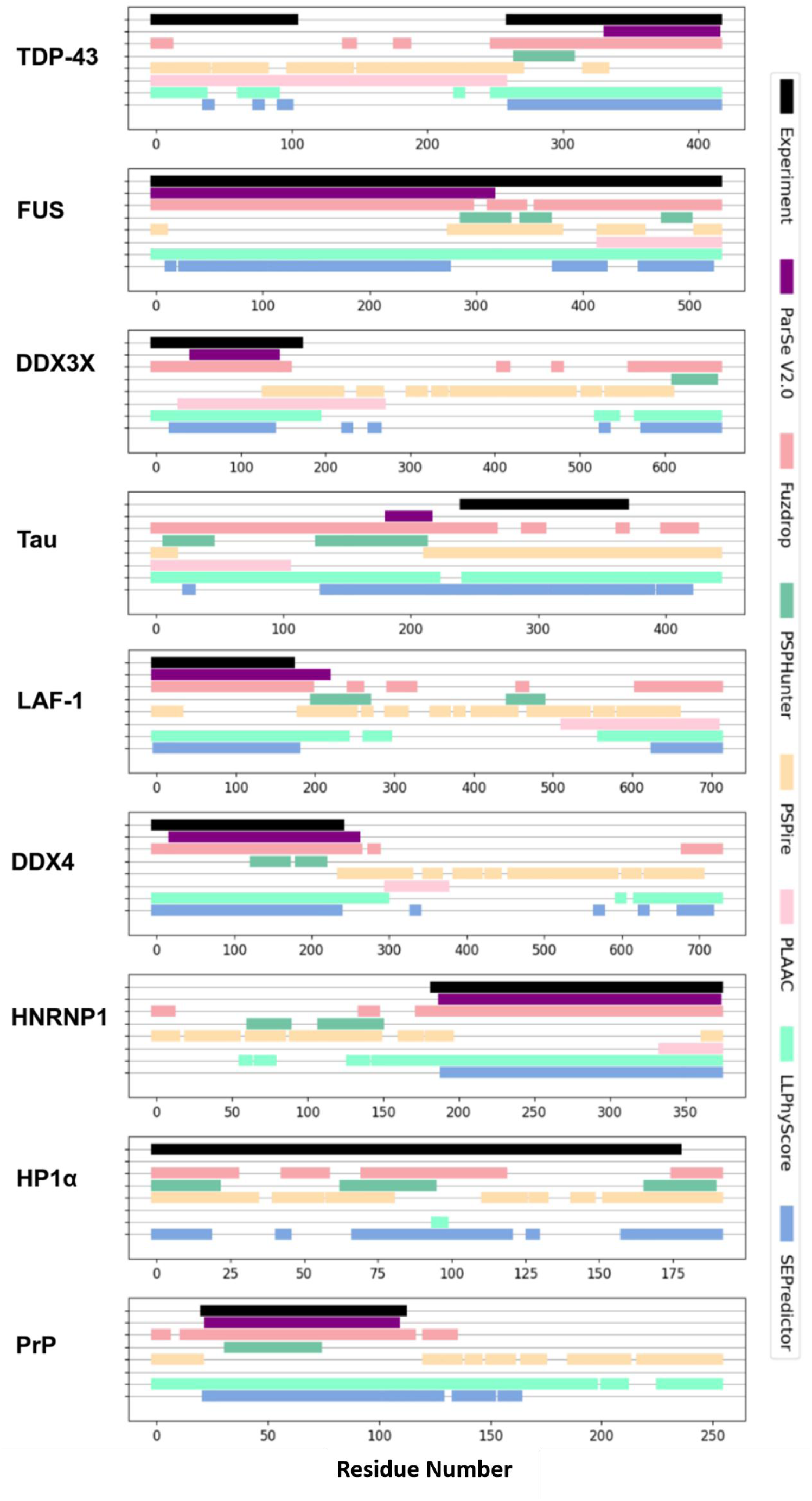
Visualization of the predicted/experimental phase-separating driving regions (based on residue scores) for representative proteins. The threshold is selected as the one corresponding to the optimal point on the ROC curve from Figure 4.

Taken together, ESM_GVP_p demonstrates superior and more robust performance across all evaluation benchmarks compared to other models developed in this study. We therefore renamed it SSPSPredictor (Structure and ESM-2-based Predictor for PSPs) and adopted it as the final model for subsequent human proteome analyses.

### IDRs more likely undergo LLPS than folded domains

To investigate the prevalence of PSPs in the human proteome and the relationship between LLPS and intrinsically disordered regions (IDRs) versus folded domains, we calculated phase separation score for each sequence of the 23,391 human proteins in AlphaFoldDB. In this study, IDR is identified as a region of continuous 30 or more amino acids with all of them having predicted local distance difference test score (pLDDT) < 70 (from AlphaFold2). Proteins lacking any IDRs were classified as “Fold proteins”. Based on whether a protein contain IDRs as well as is predicted to be a PSP, we classified human proteins in AlphaFoldDB into four categories: PSP without IDRs (Fold_LLPS), PSP with IDRs (IDRs_LLPS), non-PSP without IDRs (Fold_no_LLPS), and non-PSP with IDRs (IDRs_no_LLPS). Of the 23,391 human proteins in AlphaFoldDB, 16,668 contain IDRs, comprising 4,355 IDRs_LLPS and 12,313 IDRs_no_LLPS proteins. Meantime, the number of those belonging to Fold_LLPS and Fold_no_LLPS are 6,116 and 607 respectively. As shown in Figure 7, SSPSPredictor-based predictions indicate that approximately 10% of folded proteins undergo LLPS, compared to 35% of proteins containing IDRs—a substantially higher proportion that underscores the critical role of IDRs in phase separation. It highlights the critical role of IDRs in LLPS.

**Figure 7.**
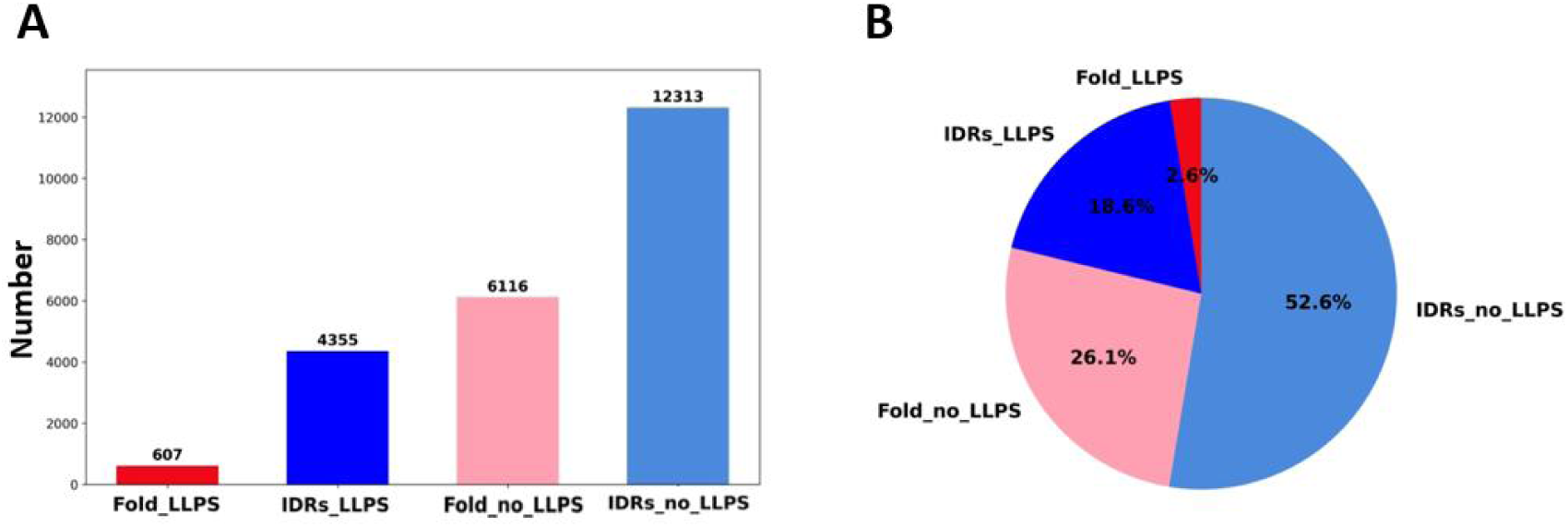
Distribution of different structural types (Fold/IDRs) of PSPs/non-PSPs predicted by SSPSPredictor for the human proteome. A: Protein numbesr. B: Protein fractions. The result is determined by the voting of the five sub-models.

### Pathogenic variants in the Human Proteome have higher attention scores

Given that the model can identify key amino acids that drive LLPS in PSPs, we explored whether pathogenic mutations are associated with phase separation. The mutation data of human proteome in AlphaFoldDB^32^ were collected from the Clinical Variation Database (ClinVar)^37^, comprising 82,643 benign and 44,256 pathogenic variants. We calculated the phase separation scores for all variant positions. Based on variant pathogenicity and local structural context—defined by pLDDT < 50 (disordered) or > 70 (folded)—we categorized all variants into four classes, including “Fold_LLPS”, “IDRs_LLPS”, “Fold_no_LLPS” and “IDRs_no_LLPS”. The statistical distribution plots of the four classes are presented in Figure 8a, which indicates that the positions of pathogenic variants usually have a higher tendency for phase separation, and this is more evident at positions within disorder regions (corresponding to low pLDDT). Figure 8b exhibits similar results using IUPred score instead of pLDDT, which is the amino-acid-level disorder score calculated by the prediction algorithm IUPred3^42^. This finding reveals that mutations within phase-separation-driving regions are more likely to be pathogenic, consistent with previous analyses^43^.

**Figure 8.**
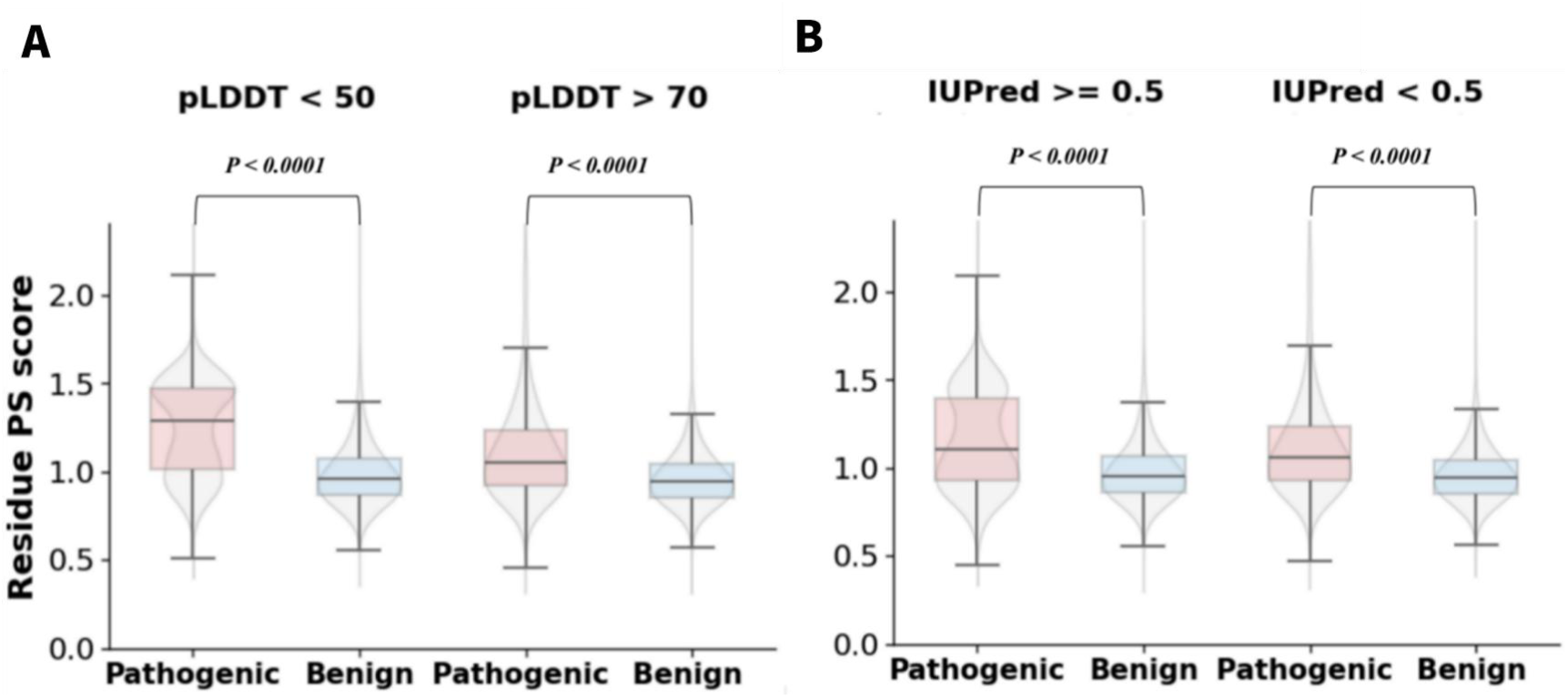
Residue-level phase-separating scores for pathogenic and benign variants on Test set 4 with different levels of disorder, predicted by pLDDT (A) and IUPred B). The result is the average of the scores from the five sub-models.

### Construction of online webserver

Finally, we developed an online server based on SSPSPredictor (http://bio-comp.ucas.ac.cn/SSPSPredictor/). Users can input either a UniProt ID (as shown in Figure 9) or a protein sequence. For a UniProt ID input, SSPSPredictor retrieves the corresponding predicted structure from AlphaFoldDB and returns the prediction rapidly. For a raw sequence input, the server first performs protein structure prediction. To accelerate structure prediction, we employed ColabFold^44^ instead of AlphaFold2, which is 40–60 times faster while maintaining comparable accuracy. When ColabFold-predicted structures were used for LLPS prediction on the test dataset, ESM_GVP_p achieved an AUROC of 0.842, AUPRC of 0.848, accuracy of 0.786, and an F1-score of 0.781—performance comparable to that obtained with AlphaFold2-predicted structures. The online prediction results include: (i) a binary classification (PSP or non-PSP), (ii) a prediction score, and (iii) a visualization highlighting the predicted crucial driving residues (if classified as a PSP). These results are automatically emailed to the user.

**Figure 9.**
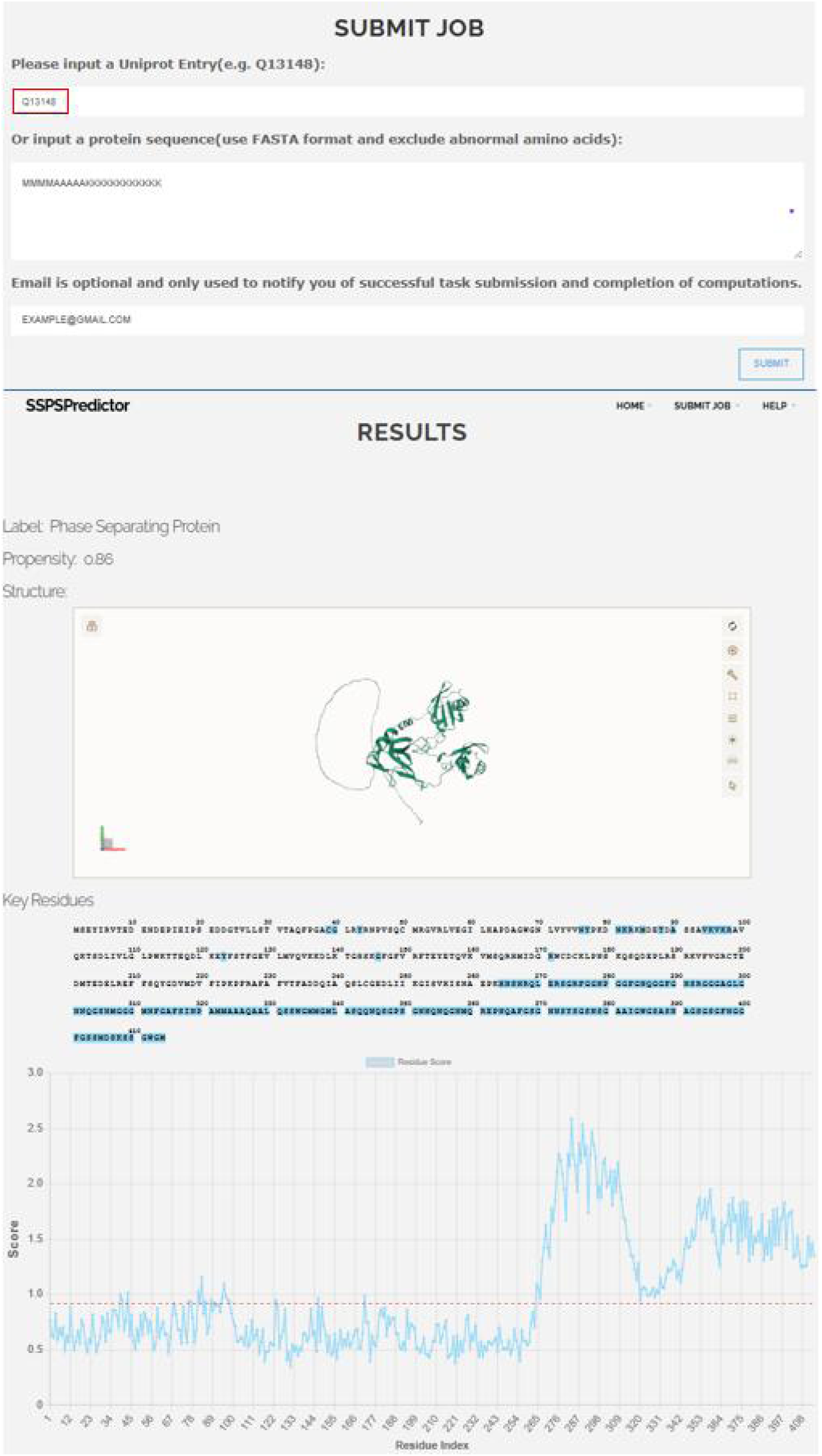
Screenshot of online server of SSPSPredictor. The schematic of prediction outcome when inputing Uniprot ID “Q13148” as an example.

## DISCUSSION AND CONCLUSION

By integrating the pre-trained protein language model ESM-2 with graph neural networks (GNNs), we developed novel models to predict phase-separating proteins (PSPs), regardless of whether they are predominantly folded or intrinsically disordered. ESM-2 automatically learns rich sequence representations, while structural information from AlphaFold2-predicted structures is integrated via graph neural networks. We evaluated two GNN architectures — GVP and SPIN-CGNN — and explored both sequential and parallel fusion strategies for combining the ESM-2 and GNN modules. The subsequent attention-pooling operation endows the model with interpretability, as evidenced by the strong correlation between its residue-level importance scores and experimentally defined phase-separation-driving regions. Comprehensive evaluation across multiple external datasets demonstrated that ESM_GVP_p—a model combining ESM-2 and GVP via parallel fusion—achieved superior and balanced performance. We therefore renamed it as SSPSPredictor (a Sequence and Structure deep learning model for Predicting Phase-Separating Proteins).

We further applied SSPSPredictor to a proteome-wide analysis of human proteins. Our analysis revealed that, contrary to the prevailing view that LLPS is primarily driven by disordered regions, a notable subset (∼ 10%) of predominantly folded proteins also exhibit phase separation propensity. Furthermore, analysis of ClinVar variants showed that pathogenic mutations are significantly enriched at residues with high LLPS propensity scores, particularly within intrinsically disordered regions. These findings highlight a strong link between dysregulated LLPS and human disease, offering a new mechanistic perspective for understanding pathogenesis.

Although SSPSPredictor represents a powerful tool for PSP prediction and analysis, several avenues for improvement remain. First, integration of emerging datasets and experimental validation—such as in vitro phase separation assays or live-cell imaging— could further refine and validate SSPSPredictor’s predictions. Second, enhancing SSPSPredictor’s scalability would enable analysis of larger and more diverse datasets, including cross-species proteomes, thereby broadening its applicability. Moreover, the model architecture could be extended to predict co-condensation propensities of protein pairs or multiprotein complexes, and—when integrated with state-of-the-art generative models—potentially guide the de novo design of synthetic PSPs.

## ACKNOWLEDGMENT

This work was financially supported by the National Natural Science Foundation of China (32571441, 32071250, 22373020) and by the Fundamental Research Funds for the Central Universities. We are grateful to Shuang Hou and Prof. Yong Zhang at Tongji University of for their assistance on PSPire, and to Dr. Pengjie Li and Dr. Yiwei Li at Huazhong University of Science and Technology for their assistance on part of test sets.

